# A benchtop bioreactor to impart coarctation-induced deformation and assess mechanisms of endothelial dysfunction

**DOI:** 10.1101/2025.05.23.655865

**Authors:** Dylan Schock, Alexander Armstrong, Abdel Alli, John F. LaDisa

**Author notes:** Address for correspondence: John F. LaDisa, Jr., PhD, 8701 W Watertown Plank Rd., Milwaukee, WI 53226.

## Abstract

Hypertension (HTN) is common with coarctation of the aorta (CoA), a common congenital cardiovascular defect. Coarctation-induced mechanical stimuli can drive remodeling in the proximal aorta, resulting in stiffening and HTN. Natriuretic peptide receptor C (NPRC) expression is downregulated in proximal aortas of hypertensive CoA patients, and preclinical models show endothelial NPRC decreases with increasing coarctation severity. The function of NPRC is regulated by actin cytoskeleton organization. AHNAK potentiates arachidonic acid calcium mobilization in this process, but its role in CoA is unknown. Mechanistic study of NPRC in CoA is hampered by limited, heterogenous patients and lengthy, expensive preclinical models. We developed an in vitro system using human aortic endothelial cells (HAECs) to mimic NPRC and endothelial dysfunction results from CoA and study mechanisms (e.g. AHNAK) regulating NPRC membrane expression. Physiologic (12%) and pathologic (17%) strain conditions derived from preclinical control and CoA aortic measurements were applied to HAECs using a tension bioreactor. Two-photon imaging of the strain-conditioned HAECs response to CNP (agonist) revealed those exposed to pathologic strain had significantly less intracellular calcium [Ca^2+^]_i_ mobilization than with physiologic strain, consistent with results from intact CoA tissue. Protein expression of NPRC and AHNAK were disrupted in actin cytoskeleton fractions from sucrose density gradient ultracentrifugation studies using lysates of HAEC exposed to pathologic conditions. These results indicate our HAECs straining system mimics the in vivo effects CoA-induced mechanical stimuli have on aortic tissue and can be used can used to conduct studies confirming and/or extending associated mechanisms.

## Introduction

Coarctation of the aorta (CoA) is a common congenital cardiovascular defect characterized by a stenosis of the proximal descending thoracic aorta^1, 2^. Surgical treatment can alleviate the stenosis but patients frequently develop cardiovascular morbidity with hypertension (HTN) being most common^3^. Studies indicate up to 80% of children with CoA will develop HTN within 25 years of surgery^4^. Difficulty unraveling the mechanisms of HTN in CoA stems from separating the causal genetic contributions putatively contributing to the initial narrowing from changes in gene expression associated with coarctation-induced mechanical stimuli imposed on the proximal aorta after closure of the ductus arteriosus.

The thoracic aorta accounts for >60% of the total arterial compliance. Hence, adverse coarctation-induced mechanical stimuli in this region can drive remodeling resulting in aortic stiffening and HTN^5^. Computational simulations using a preclinical (rabbit) CoA model confirmed the mechanical stimuli for remodeling above CoA are rooted in pronounced deformation (quantified as strain)^6^ from elevated blood pressure (BP). Specifically, chronic changes in wall tension (radius x BP) experienced by arterial cells is believed to be the stimuli for thickening via remodeling. While thickening restores wall stress to a preferred range^7^, it adds stiffness thereby decreasing strain. Decreased strain occurs with age^8^, is a manifestation of arterial remodeling, and is an indicator of HTN risk^9, 10^. Importantly, recent work^11, 12^ has determined the range of values for BP gradients, walls stress and strain leading to persistent arterial dysfunction in response to the severity and duration of CoA in the same preclinical model referenced above using data that are also routinely obtained as part of the standard of care for patients treated for CoA.

We previously identified a candidate gene from patients with CoA having upper extremity systolic BP >99th percentile for their gender, height and age, thereby presumably focusing on mechanical stimuli from the coarctation^13^. In this work, RNA sequencing (RNAseq) of tissue from CoA patients revealed downregulation of *natriuretic peptide receptor C (NPRC;* a.k.a. *NPR3)* in proximal thoracic aorta sections subjected to high BP when compared to distal sections exposed to normal BP. Importantly, microarray data from the preclinical CoA model referenced above^6, 14^ also showed downregulation of *NPRC* in proximal aortas from both untreated and treated CoA rabbits experiencing localized mechanical stimuli, including elevated BP and pronounced deformation, as compared to controls^11^. Aortic tissue from the high BP region also showed impaired endothelial function to acetylcholine, even in animals where the BP was restored to normal for the equivalent of 6 human years^15^. Smooth muscle (SM) relaxation was normal in aortic tissue from the high BP region despite persistent medial thickening and stiffening^6^. The functional relevance of coarctation-induced changes in *NPRC* manifested as impaired relaxation and intracellular calcium transients [Ca^2+^]_i_ from aortic tissue samples taken proximal to the coarctation in this model^13^.

NPRC is one of 3 natriuretic peptide receptors that include atrial (ANP), brain (BNP) and C-type (CNP). ANP and BNP are mostly found in the atria and ventricles, whereas CNP is abundant in vascular EC^16, 17^. ANP, BNP and CNP all have strong affinity for NPRC^18^, a transmembrane domain receptor coupled to adenylyl cyclase (AC) inhibition via inhibitory guanine nucleotide regulatory protein (Gi)^19^. Natriuretic peptides favorably impact cardiovascular function. For example, their impairment is implicated in type 2 diabetes, obesity and heart failure^20-22^. Specifically, NPRC is important for its role in regulating systemic BP^19^ via AC inhibition^23^. NPRC is found in tissues including endothelial cells (EC) where it plays a role in re-endothelialization and viability under healthy conditions^24^. We have previously shown^13^ relaxation of the proximal aorta to CNP and ANP is abolished by pre-treatment with the NPRC inhibitor^25^, M372049.

There are 2 subtypes of NPRC with molecular masses of 67-kDa and 77-kDa that are physiologically relevant to CoA. The 77-kDa protein is involved in ligand binding and functions mainly as a clearance receptor to remove circulating natriuretic peptides^26^. The 67-kDa protein is linked to activation of phospholipase C (PLC) and inhibition of AC NPRC is also coupled to endothelial nitric oxide synthase (eNOS) activity required for nitric oxide (NO) release^27^.

Importantly, endothelial sodium channel (EnNaC) activity results in EC stiffening and suppression of NO production^28^, while absence of EnNaC results in “softening” of EC and production of NO^28-33^. Increased BP is also linked to decreases in EnNaC activity via feedback inhibition^34^. Anionic phospholipid phosphates such as phosphatidylinositol (PI)-4,5-bisphosphate are important intracellular signal transducers that positively regulate channel activity by maintaining an open conformation^35-38^. AHNAK is a large multifunctional scaffolding protein that is known to regulate the cytoarchitecture of the cell membrane by interacting with the annexin 2/S100A10 complex^39^. The carboxyl terminal domain of AHNAK has been reported to serve as an adaptor between L-type calcium channels and the actin cytoskeleton^40^. AHNAK was also shown to interact with NPRC at the plasma membrane to potentiate arachidonic acid mediated calcium mobilization^41^. Collectively these prior studies suggest AHNAK may play a role deformation-induced cellular stiffening, but its role in CoA has not been studied to date.

Further studying the involvement of this promising NPRC pathway relative to coarctation-induced deformation and HTN in patients is intractable given the relatively low number of CoA cases at each institution annually and their heterogeneity since CoA can present with many conditions including bicuspid aortic valve^4^. The findings reviewed above were obtained from commercially available aortic EC lines subjected to methods that agonize and/or inhibit NPRC and its associated pathway, but associated mechanisms have not yet been fully elucidated in the setting of coarctation-induced deformation. Beyond being representative of the clinical condition, any cellular-based system designed to scrutinize associated mechanisms should also be efficient, since preclinical models can be expensive and the use of agonists/inhibitors with harvested tissue or in vivo may not yield physiologic relevance.

The goals of the current study were to (1) develop an in vitro system using cultured ECs to mimic and complement NPRC and endothelial dysfunction results from our preclinical model of CoA, (2) use this system to begin further elucidating associated mechanisms of EC dysfunction, and (3) investigate for the first time the regulation of AHNAK and NPRC proteins in HAEC in response to pathological strain.

Specifically, we used a tension bioreactor to apply physiologic control or pathologic CoA strain conditions measured from the preclinical model to cultured human aortic ECs (HAEC).

Two-photon imaging was used to quantify the strain-conditioned [Ca^2+^]_i_ response when interrogated with CNP and sucrose density gradient ultracentrifugation studies were then conducted based on our collective knowledge and convergence of pathways related to *NPRC* and coarctation-induced deformation. Our hypothesis was that applying pathologic CoA levels of strain to cultured HAECs would cause a less pronounced [Ca^2+^]_i_ response to a dosage of CNP as compared to HAECs that underwent physiologic control levels of strain, consistent with prior results from intact aortic tissue^13^. Our results suggest the system and protocol developed permits investigation of the mechanisms associated with deformation-induced downregulation of NPRC and its potential application to HTN, including AHNAK.

## Materials and Methods

### Primary human aortic endothelial cells

HAECs from Cell Biologics (Chicago, IL) were cultured and expanded using the manufacturers recommended human endothelial cell medium supplemented with 5% fetal bovine serum, 10mL/L of L-Glutamine and Antibiotic-Antimycotic, and 1mL/L of VEGF, HEPARIN, EGF, and FGF. Cultures were maintained in a humidified incubator at 37°C and an atmosphere of 5% CO2-95% air. Cells were received frozen at passage 3 and initially seeded onto a T25 cell culture flask or stored in liquid nitrogen. Cell media was replenished every 2 days and passaged in a 1:3 ratio every 5-6 days. Cell viability was monitored daily by performing qualitative checks of morphology, pH indicator color, and visual inspection for bacterial/fungal contamination. All culture flasks were coated with a Gelatin-Based Coating Solution (Cell Biologics) for at least 2 minutes and then aspirated prior to seeding. Cells were split in a 1:2 ratio onto T75 culture flasks at passage 7. Once >80% confluent, cells were either frozen and stored at -80°C for later experiments or seeded at 100,000 cells/well onto a six-well Bioflex plate coated with Collagen I from FlexCell International Corporation (Burlington, NC). Experimental cell populations were typically seeded about 2 days before any planned experiment to ensure confluency. Experiments were conducted at passage 8 (P8) to maximize the experimental population and cell viability while also aligning with methods and results from historically published data^13^.

### Imposing CoA-induced deformation via bioreactor mimics impaired [Ca^2+^]_i_ transients from intact aorta

To replicate deformation-induced changes in NPRC governing dysfunction, the cultured HAECs mentioned above were exposed to cyclic equibiaxial strain values measured from control (12%; normal) and CoA (17%; pathologic) groups of rabbits prior to the onset of remodeling using phase-contrast MRI^6, 42^. The differences in strain were imposed on EC using via bioreactor (FX-6000T; FlexCell Int). Strain was set to be applied cyclically for one hour using a triangular waveform. A duty cycle of 33% was similarly set such that 1/3 of the waveform is spent increasing in strain and 2/3 is spent decreasing back to a baseline of 0% elongation. This pattern was chosen to mimic the displacement pattern experienced by the aortic wall over the course of a cardiac cycle^43, 44^.

CNP-induced hyperpolarization requires Ca^2+^ from intracellular stores or the extracellular space^45^. Average [Ca^2+^]_i_ mobilization was therefore recorded (n=4-6 wells/plate) during experimentation that followed cyclic straining of cells. Imaging was performed using an upright Fluoview FV1000 Laser Scanning 2-photon (aka multiphoton) microscope from Olympus (Tokyo, JP) equipped with Ti:sapphire lasers set to a wavelength of 820nm. Samples were imaged using a 25X (N.A. 1.05 and working distance 2mm) water-immersion objective lens (XLPL25XWMP, Olympus). Prior to imaging, the strain-conditioned HAECs were loaded with the Ca^2+^ sensitive dye Fluo-4 AM (3.8μM, Invitrogen) in dimethyl sulfoxide (DMSO) using 0.02% Pluronic acid (Pluronic F-127, Invitrogen) in basal media for one hour. The flexible silicone Bioflex membranes containing the cells were carefully sectioned from the Bioflex plate using a scalpel and pinned down to a silicone elastomer-coated 35mm imaging dish. The membranes were immersed in warmed 2mM Ca^2+^ physiologic salt solution (PSS) consisting of Millipore-filtered water and 8.47g/L NaCl, 2.60g/L HEPES, 0.222g/L CaCl2, 0.190g/L MgCL2, and 0.335g/L of KCl.

Once the silicone membranes containing strain-conditioned ECs were securely pinned down in the tissue culture dish, the dish was transferred into the 2-photon apparatus. Brightfield imaging was used to locate the EC layer and the microscope was then switched into multiphoton mode. Continuous digital image acquisition settings were set to capture 16-bit images every 1.644s for a maximum duration of 200 frames (328.5s). After starting the experiment, 30 frames were allowed to pass to establish a baseline intensity reading of the cells prior to the addition of any reagents or peptides. At frame 30, 10mL of warmed PSS containing 1.5μM of CNP was gently diluted into the imaging dish via syringe while the same volume of PSS in the dish was simultaneously removed. This dose was chosen to balance cellular response to the agent with toxicity and was further scrutinized in separate dose dependance studies (0.5, 1.5 or 5 uM)^13, 46^. Digital images of each frame during experiments were saved for offline analysis. Changes in intensity reflective of [Ca^2+^]_i_ mobilization were imaged with this protocol using ECs subjected to the physiologic control or pathologic CoA strain conditions mentioned above. Biological replicates (n = 3 Bioflex plates) were obtained using measurements from at least 3 experiments in most cases, with each plate included in the study containing at least one technical replicate (n ≥ 3 wells).

Bioreactor results were characterized for EC in response to a deformation protocol (**Fig 1**) when NRPC activity was inhibited or augmented by multiple approaches. More specifically, [Ca^2+^]_i_ intensity was measured in ECs exposed to physiologic control or pathologic CoA strain conditions and then treatment with the NPRC-specific pharmacological inhibitor, M372049 (AstraZeneca)^25^. The deformation duration used is supported by studies showing 1-hour of cyclic strain on HAEC is sufficient to induce changes in gene expression and phenotype^47-49^. Additional tests were conducted for the current study to determine that [Ca^2+^]_i_ differences between physiologic control or pathologic CoA straining were maintained after 12 hrs of cyclic strain regardless of quantification applied (see Quantification below).

**Fig 1.**
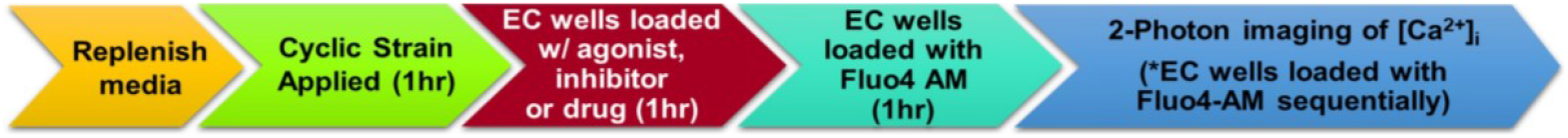
Protocol to study EC NPRC after CoA-induced deformation.

### Quantification

Images were quantified using the Loci tool plugin in Fiji (aka ImageJ; National Institute of Health). Regions of interest (ROIs) were identified over five different cells at random, as well as an area of empty space to represent the background fluorescent intensity. The Measurement option within the Analyze tool was employed to record the mean intensity, among other features related to fluorescent intensity across all frames. Temporal measurements for each of the 5 recorded ROIs and background were cataloged and saved. The raw mean data from each ROI was loaded into Origin6.0 (OriginLab), where the background intensity values for each frame were subtracted from the 5 ROIs to normalize results. The mean Fluo4-AM intensity values (now background corrected) for the first 30 frames before CNP application were averaged for each ROI, creating a baseline value to compare the subsequent change in [Ca^2+^]_i_. percent change in the [Ca^2+^]_i_ relative to the baseline for each recorded time point was calculated.

The resulting changes in percent [Ca^2+^]_i_ for each ROI were then graphed and a moving average baseline was calculated from the first ∼30 frames prior to CNP application using Origin6.0. The temporal data from each biologic replicates for each condition processed were then loaded into MATLAB for further quantifications and statistical analysis. Average [Ca^2+^]_i_ mobilization and standard error of the mean over time were calculated for each well and plate. Quantifications were conducted consistent with methods applied for intact arterial segments from a preclinical rabbit model of CoA^13^. Specifically, the temporal [Ca^2+^]_i_ mobilization curves and initial absolute decreases in [Ca^2+^]_i_ after CNP application were quantified and graphed to compare with results from the in vivo rabbit model. Additionally, the area under the curve (AUC) values for each of the temporal [Ca^2+^]_i_ graphs were quantified by calculating the definite integral of the curves from 0 to 150 seconds. One-way analysis of variance (ANOVA) was performed between each condition to determine statistical significance defined as samples having a *P*-value ≤ 0.05. Prism 9 (GraphPad) was used to test for statistical differences and to create associated figures.

### Sucrose Density Gradient Ultracentrifugation Assays and Western blotting

Lysates from HAEC subject to physiological or pathological strain were subject to sucrose density gradient ultracentrifugation as described by Montgomery et al^50^ with the following modifications. HAEC were scraped in lysis buffer and then mixed in an equal volume with freshly prepared ice-cold 1% Brij 96/TNEV buffer (10 mM Tris·HCl, pH 7.5, 150 mM NaCl, 5 mM EDTA, 2 mM sodium vanadate. The suspension was passed 15 times through a 23-gauge syringe before being incubated on ice for 1 h. Next, the lysates were subject to centrifugation at 10,000 rpm at 4°C for 5 min to remove cellular debris, and 500 µl of the supernatant were mixed with 500 µl of freshly prepared 80% sucrose in TNE (without Brij 96 and without vanadate) before being transferred to a 13 × 23-mm Beckman centrifuge tube. Next, 1,800µl of 35% sucrose in TNE was carefully applied to the top of the mixture followed by application of 500 µl of 5% sucrose. The sucrose gradient was subject to ultracentrifugtion at 34,000 rpm at 4°C for 16 h in a SW50.1 rotor (Beckman). Finally, fourteen fractions of 235-µl volumes were collected from the top to the bottom of the tube being careful not to disturb the adjacent fractions. The other fractions were resolved by SDS-PAGE and probed for ANHAK and NPRC proteins by Western blotting. Rabbit polyclonal AHNAK1 antibody was provided by Dr. Hannelore Haase (Max Delbrück Center, Berlin, Germany).

## Results

Initial analysis of trends from HAECs exposed to the experimental protocol above (**Figure 2**) indicated CNP-induced [Ca^2+^]_i_ changes consistent with intact aorta exposed to CoA^13^, confirming the utility of our cyclic straining bioreactor to uncover NPRC-dependent mechanisms of dysfunction from excessive EC deformation. Additional trials were therefore conducted using the protocol and associated methods.

**Fig 2.**
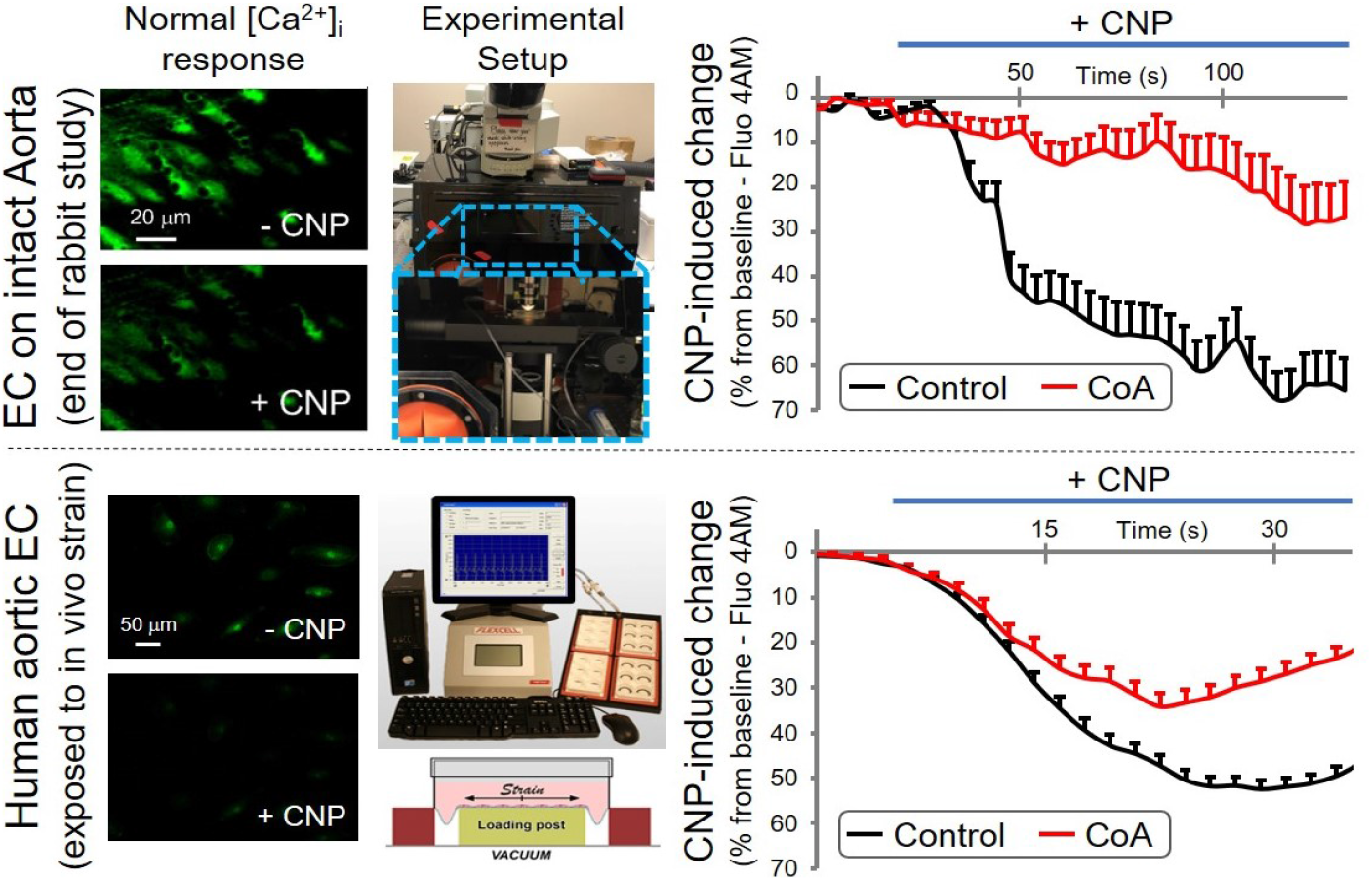
CNP-induced changes in [Ca^2+^]_i_ confirm our cyclic straining that exposes EC to excessive deformation from CoA before remodeling (bottom) mimics results from intact aortic segments (top).

### Physiologic Control vs. Pathologic CoA Strain

All groups of strain-conditioned HAECs tested generally showed a rapid decrease in [Ca^2+^]_i_ (% relative to baseline) when exposed to a bolus of 1.5μM of CNP. HAECs exposed to normal levels of cyclic strain (i.e., 12%; Physiologic Control group) for 1 hour had a more pronounced reduction in [Ca^2+^]_i_ than ECs exposed to pathologic cyclic strain (i.e. 17%; Pathologic CoA group) for the same duration. This behavior can be seen in **Figure 3A left**, where the mean temporal [Ca^2+^]_i_ was tracked for 100 seconds after administration of CNP (∼50 seconds after the start of imaging). Both groups of HAECs had initial decreases in [Ca^2+^]_i_, but to differing levels, followed by a partial return towards their pre-CNP baselines after dilution. The partial recovery of the lost cytosolic [Ca^2+^]_i_ peaked between 55 and 70 seconds after CNP.

**Fig 3.**
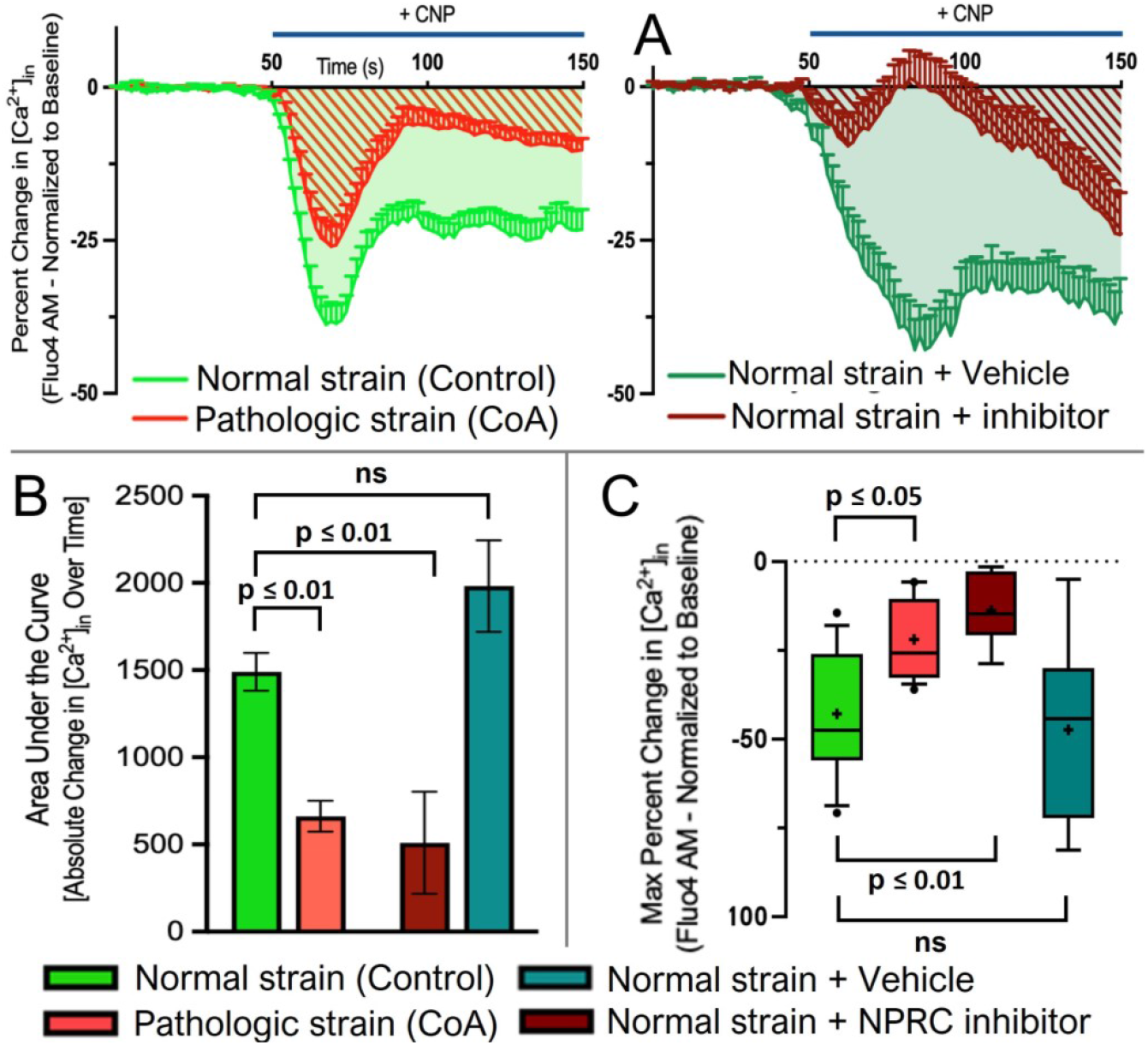
(A) Data of [Ca^2+^]_i_ from CNP depending on severity of strain and NPRC inhibition (10 μM M372049 for 60 min). (B&C) [Ca^2+^]_i_ quantification methods (n=3-6 wells/group).

To further confirm a role for NPRC in the current study, [Ca^2+^]_i_ intensity was measured in HAECs exposed to physiologic strain (12% elongation) conditions and then treated with the NPRC-specific pharmacological inhibitor, M372049 (AstraZeneca)^25^. HAECs that underwent 1 hour of physiologic cyclic strain followed by incubation with the NPRC-specific inhibitor (i.e., Normal strain + Inhibitor) had an impaired response to CNP as compared to incubation with the vehicle, DMSO (i.e., Normal strain + Vehicle), as shown in **Figure 3A right.** These patterns generally mimic the HAEC responses to CNP after undergoing Physiologic Control vs Pathologic CoA strain levels. Following the initial CNP-induced decrease in [Ca^2+^]_i_, the Physiologic + Vehicle group did exhibit partial [Ca^2+^]_i_ recovery, while the Physiologic + Inhibitor cells had increased cytosolic [Ca^2+^]_i_ levels that went beyond their pre-CNP baselines briefly before steadily decreasing from the initial baseline.

The AUC of the absolute change in [Ca^2+^]_i_ from 0 to 150 seconds was quantified **Figure 3B** using the definite integrals of the [Ca^2+^]_i_ responses (Figure 4-right, shaded regions) and compared across experimental groups. The physiologic control strain group had a higher mean AUC (1490 ± 108 [Ca^2+^]_i_ %*s) than the pathologic CoA strain group (662 ± 89 [Ca^2+^]_i_ %*s, p ≤ 0.01) and Normal strain + Inhibitor groups (510 ± 292 [Ca^2+^]_i_ %*s, p ≤ 0.01). The AUC of the Normal strain + Vehicle group (1980 ± 263 [Ca^2+^]_i_ %*s) was not significantly different from the Normal strain Control group.

**Fig 4.**
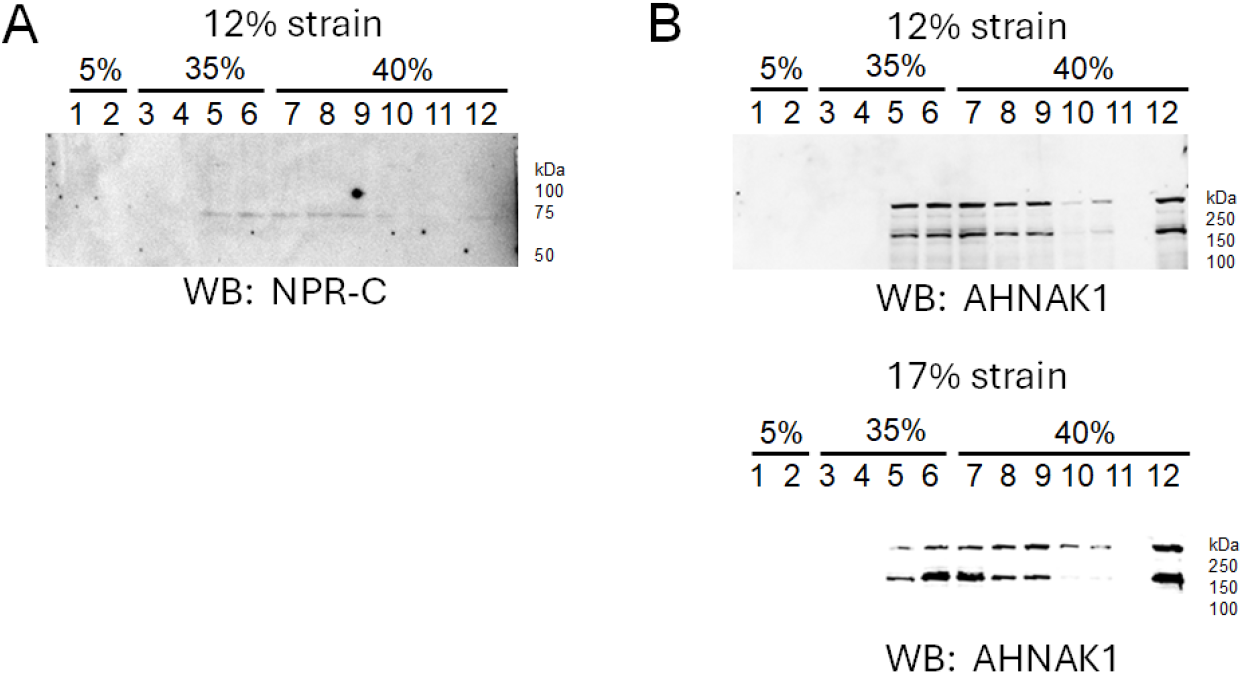
Sucrose Density Gradient Assay and Western blot analysis of AHNAK1 protein expression in different fractions from HAEC subject to physiologic or pathologic strain. (A) Western blot of NPRC protein expression in medium density fractions from HAEC subject to physiologic control stain, (B) Western blot of AHNAK1 protein expression predominantly in medium density fractions from HAEC subject to physiologic strain (top blot), (C) Western blot of AHNAK1 protein expression dispersed in medium and heavy density fractions from HAEC subject to pathologic strain. In both physiologic strain and pathologic strain there is some AHNAK1 expressed in the nuclear fraction (last fraction).

The maximum percent decrease in cytosolic [Ca^2+^]_i_ after CNP administration was extracted for all conditions. **Figure 3C** shows the Normal strain group had a significantly more pronounced [Ca^2+^]_i_ decrease in response to CNP (−42.9 ± 2.1%) as compared to the Pathologic CoA (−22.0 ± 2.6%, p ≤ 0.05) and Normal strain + Inhibitor groups (−13.8 ± 2.9%, p ≤ 0.01). The maximum percent decrease of [Ca^2+^]_i_ in the Normal strain + Vehicle group was not significantly different (−47.4 ± 4.2%) from the Normal strain group.

### AHNAK1 protein expression shifts from lighter density to heavy density sucrose gradient fractions in HAEC subjected to pathological strain

A previous study showed the AHNAK1 C1 domain interacts with the cytoplasmic tail of NPRC and siRNA mediated knockdown of NPRC results in AHNAK1 protein expression shifting from cytoplasmic and membrane fractions to nuclear fractions in AoSMC’s^41^. Here we investigated whether pathological strain in HAEC alters AHNAK1 membrane expression compared to cells exposed to normal strain. As shown in **Figure 4**, AHNAK1 is shown to shift from medium to heavy density gradient fractions in response to pathologic strain.

## Discussion

A critical barrier to our understanding of HTN in CoA involves separating causal genetic contributions from changes in gene expression due to mechanical stimuli after closure of the ductus arteriosus. We previously identified *NPRC* as a candidate gene from CoA patients with upper extremity systolic BP >99th percentile for their sex, height and age, thereby focusing on mechanical consequences^13^. RNA sequencing showed downregulation of *NPRC* in proximal aortic sections subjected to high BP versus distal sections exposed to normal BP. Microarray data from a CoA animal (rabbit) model^6, 14^ also showed downregulation of *NPRC* in proximal aortas from CoA and treated animals experiencing high BP and pronounced deformation. Aortic tissue from the high BP region also revealed impaired endothelial function to acetylcholine and natriuretic peptide agonists despite restoration of normal BP for the equivalent of 6 human years^15^. SM relaxation is unchanged in these models despite medial thickening and stiffening^6^, which are normal adaptive responses to elevated wall tension^51^. These data strongly suggest endothelial NPRC is a key component of the post-CoA HTN mechanism, yet to our knowledge, there are limited studies of EC NPRC activity from pronounced deformation (e.g. CoA).

Our research to uncover the link between mechanical stimuli and HTN in CoA has led to the need for a cellular-based system to scrutinize associated mechanisms in a manner that is representative of the clinical condition and also efficient. Mechanistic study in humans is intractable given the relatively low number of CoA patients at each institution and their heterogeneity. Prior efforts using our preclinical model to identify mechanistic targets have been fruitful^13, 52^, but are slow and resource limiting as these preclinical models can be expensive. For example, our early work identified mechanistic targets (e.g. SERCA)^52^ for which follow-up studies using arteries harvested after ∼6 month experiments with associated agonists and inhibitors have not revealed changes in active force assessment via myography to date. Hence, the goal of the current study was to develop a rapid approach to study mechanisms of deformation induced changes in endothelial NPRC. Promising mechanistic candidates could then be used in follow-up preclinical studies to confirm physiologic relevance in vivo pending favorable results.

There are 3 main findings from the current study. (1) CNP-induced [Ca^2+^]_i_ changes measured in HAEC of the current study after normal and pathologic straining are consistent with control intact aorta and that exposed to pronounced deformation from CoA^13^, (2) pharmacological impairment using the NPRC-specific inhibitor, M372049, produced [Ca^2+^]_i_ transients similar to those seen after exposure of HAEC to pathologic levels of strain mimicking CoA, (3) AHNAK seems to play a role in the mechanisms of these NPRC changes. Each of these findings is discussed in more detail below.

The current finding of CNP-induced [Ca^2+^]_i_ changes to CNP following control or pathologic levels of strain being consistent with results from intact aorta exposed to CoA^13^ provided confidence that our cyclic straining bioreactor could be used to uncover NPRC-dependent mechanisms of dysfunction from excessive EC deformation. Associated mechanism within the results and described below will be further assessed with our preclinical models and CoA patient tissue to further scrutinize or iterate on this bioreactor approach.

The changes observed in response to CNP occurring after 1 hour of cyclic strain applied to cultured HAECs are aligned with other studies. Multiple studies have shown that 1-hour of cyclic stretch on aortic ECs is sufficient to induce physiologic changes in both gene expression and phenotype, some of which can influence the expression of NPRC^47-49^. For example, it has been shown that Angiotensin II, which increases in ECs after 1 hour of cyclic strain, decreases the concentration of *NPRC* mRNA in vascular SM cells^53^. While the gene expression for *NPRC* was decreased in this study, the expression of *NPRA* and *NPRB* was not significantly altered.

Bovine aortic ECs were found to have an increase in adenylyl cyclase activity concentration of cAMP after 1 hour of cyclic strain. Increases in both cytosolic cAMP and cGMP concentration have been shown to inhibit *NPRC* gene expression^54^. Increased cGMP concentrations have been shown to inhibit PLC signaling activity, which in turn decreases the production of DAG and IP; both of which are generated by phosphatidyl inositol bisphosphate and involved with [Ca^2+^]_i_ signaling and activation of PKC respectively^19^. PKC activity has been shown to be increased in cultured aortic ECs after 1 hour, as HeLa cells that have had increased PKC activation have also been shown to demonstrate downregulated NPRC expression through both transcriptional and post-transcriptional mechanisms after applying strain for up to 1 hour^55^. Interestingly, PKC activity in aortic ECs exposed to cyclic strain is significantly elevated after just 5 minutes of cyclic stretch. This upregulated PKC state starts to steadily decrease until 1 hour of strain is applied, where the upregulation then becomes static. This behavior indicates that there are substantial physiologic changes taking place within the first 1 hour of cyclic strain, with diminishing returns for strain that is applied beyond 1 hour. PKC is a primary target of DAG and regulates many intracellular responses, however the specific molecular mechanisms associated with this interaction are not fully understood. Recent studies have focused on the impact of protein kinase D (PKD), which has known implications in cell growth, gene expression, and protein trafficking in ECs. DAG regulates both the localization of PKD and phosphorylates PKD by using PKC^56^.

Although membrane-cytoskeleton interactions are known to play an important role in regulating the function of transmembrane receptors^57^, the regulation of these interactions in the context of pathological strain is largely unknown. Here we investigated changes in protein expression of the cytoskeleton scaffolding protein AHNAK1 and NPRC in HAEC subjected to pathologic compared to physiologic strain. Since a previous study by Alli and Gower identified a direct interaction between NPRC and AHNAK1 in aortic vascular smooth muscle cells, rat gastric mucosa 1 cells, and 3T3-L1 adipocytes^41^, it was not surprising to observe AHNAK1 colocalizing with NPRC in HAEC. Importantly, HAEC subject to pathologic strain resulted in AHNAK1 protein expression shifting from medium density fractions to heavy density gradient fractions. This indicates there is less association between AHNAK1 with NPRC and presumably translocation of the protein to the nucleus during pathologic strain.

A link between HTN and endothelial dysfunction is well documented^58, 59^, such as impaired endothelial function in Framingham Heart Study offspring correlating with HTN severity^60^. Whether dysfunction is the cause or effect of HTN is unknown^61^, with recent studies suggesting pronounced deformation *precedes* endothelial dysfunction^62, 63^. Our data from a preclinical CoA model show NPRC expression and endothelial function are increasingly impaired with CoA severity^12^. Mechano-transduction between endothelial and medial layers is known to occur via players (e.g. stretch activated channel and integrin proteins), some of which are linked to our novel NPRC-based mechanism. The current study therefore imposed control and pathologic levels of strain informed from in vivo data to uncover the mechanisms of EC dysfunction impacting NPRC.

The current results should be interpreted within the constraints of several potential limitations. We did not study wall shear stress indices mainly because they contribute distal to CoA, with almost no differences proximally.^6,10^ EC results from our prior preclinical CoA model were contained within intact aortic segments. This is in juxtaposition to the ECs in the current study that are of human origin and were grown in culture. HAECs represent a single cell type growing in a 2-dimensional monoculture environment free of the systemic changes that are found within the animal during homeostasis and experimentation, along with the influence of mechanical stimuli mentioned before. The flexible membranes in the Bioflex plates upon which HAECs were cultured are capable of applying uniform radial and circumferential cyclic strain. The current study aimed to match the radial strain values derived from our rabbit CoA model prior to the onset of remodeling, but these values undoubtedly change as the aorta remodels in response to the severity and duration of CoA^11, 12^.

The 1-hour study duration applied in the current investigation is favorable as it limits the potential for failed experiments while allowing for the further assessment of NPRC-induced mechanisms associated with elevated levels of strain. Nonetheless, the impact of the length of time that cyclic strain was applied in vitro on HAEC function was further evaluated by comparing the [Ca^2+^]_i_ responses to CNP after undergoing 12 hours of cyclic strain in separate groups of HAEC as discussed in detail elsewhere^64^. Even though pathologic strain has a more pronounced effect on cell function after 12 hours, the difference between the physiologic and pathologic strained HAECs remained significant despite whether strain is applied for 1 or 12 hours. Thus, 1 hour of applied cyclic strain is sufficient to induce a significant difference in the CNP-induced [Ca^2+^]_i_ mobilization between control and pathologic conditions. Similarly, When comparing the effects of the various CNP concentrations directly (0.5, 1.5 and 5 uM CNP) as described elsewhere^64^, there were significant differences between the physiologic and pathologic strain groups when treated with either 1.5μM or 5μM of CNP. In contrast, 0.5μM of CNP was not potent enough to induce a significantly different [Ca^2+^]_i_ in response between the strain conditions when using the quantification methods described above.

## Conclusion

Collectively, the current results show the in vitro model developed using ECs in a tension bioreactor successfully mimicked the in vivo effects that CoA-induced mechanical stimuli have on the aortic tissue from the rabbit model^13^ in that the [Ca^2+^]_i_ response to CNP in strain-conditioned HAECs was generally consistent with the data observed from interrogating ECs from intact aortic segments. Therefore, this model can reasonably be used to help unravel the mechanisms behind coarctation-induced downregulation of NPRC (e.g. AHNAK) and their potential involvement in the development of chronic HTN. Future investigations into these underlying mechanisms can potentially provide the framework to discover underlying new therapies for CoA-induced endothelial dysfunction and HTN. The current system may therefore be used as a tool to screen possible NPRC agents, agonist and inhibitors prior to their use to reduce HTN in preclinical models and/or patients with CoA.

## Acknowledgements

The authors acknowledge the mentorship of Brandon Tefft PhD and Joy Lincoln PhD throughout this study, as well as the technical assistance of Marie Schulte PhD, Oleg Palygin PhD, and Hilda Jurkiewicz.

